# Corner flows induced by surfactant-producing bacteria *Bacillus subtilis* and *Pseudomonas fluorescens*

**DOI:** 10.1101/2022.06.20.496927

**Authors:** Yuan Li, Joe Sanfilippo, Daniel Kearns, Judy Q. Yang

**Affiliations:** Saint Anthony Falls Laboratory, University of Minnesota, Minneapolis, Minnesota, USA; Department of Civil, Environmental, and Geo-Engineering, University of Minnesota, Minneapolis, Minnesota, USA; Department of Biochemistry, University of Illinois at Urbana–Champaign, Urbana, Illinois, USA; Department of Biology, Indiana University, Bloomington, Indiana, USA

**Author notes:** Address correspondence to Judy Yang,. Judy Yang and Yuan Li conceived the idea, designed the research, and wrote the paper. Yuan Li designed and conducted the experiments. Joe Sanfilippo and Daniel Kearns contributed strains and contributed to the research idea and writing.

## Abstract

Mechanistic understanding of bacterial spreading in soil is critical to control pathogenic contamination of groundwater and soil as well as design bioremediation projects. However, our understanding is currently limited by the lack of direct bacterial imaging in soil conditions. Here, we overcome this limitation by directly observing the spread of bacterial solution in a transparent chamber with varying corner angles designed to replicate soil-like conditions. We show that two common soil bacteria, *Bacillus subtilis* and *Pseudomonas fluorescens*, generate flows along sharp corners (< 60°) by producing surfactants that turn nonwetting solid surfaces into wetting surfaces. We further show that a surfactant-deficient mutant of *B. subtilis* cannot generate corner flows along sharp corners, confirming that the bacteria-generated corner flows require the production of bacterial surfactants. The speed of biosurfactant-induced corner flow at the sharp corner is about several millimeters per hour, similar to that of bacterial swarming, the fastest mode of known bacterial surface translocation. We further demonstrate that the bacteria-generated corner flow only occurs when the corner angle is less than a critical value, which can be predicted from the contact angle of the bacterial solution. Furthermore, we show that the corner flow has a maximum height due to the roundness or cutoff of corners. The mechanistic understanding and mathematical theories of bacterial spreading presented in this study will help improve predictions of bacterial spreading in soil, where corners are ubiquitous, and facilitate future designs of soil contamination mitigation and other bioremediation projects.

**Importance:** The spread of bacterial cells in soil regulates soil biogeochemical cycles, increases the possibility of soil and groundwater contamination, and controls the efficiency of many bacteria-based bioremediation projects. However, mechanistic understanding of bacterial spreading in soil remains incomplete due to a lack of direct or in-situ observations. Here, we simulate confined spaces of soil using a transparent material with similar hydrophobicity as hydrocarbon-covered soil and directly visualize the spread of two common soil bacteria, *Bacillus subtilis* and *Pseudomonas fluorescens*. We show that both bacteria can generate vertical flows along sharp corners of the transparent chamber. The velocity of the bacterial corner flow is several millimeters per hour. We further demonstrate that the corner flow was generated by bacteria-produced bio-surfactants, which are soap-like chemicals and turn nonwetting solid surfaces into wetting surfaces. Our results will help improve predictions of bacterial spreading in soil and facilitate designs of soil-related bioremediation projects.

## Introduction

Bacteria play a major role in soil carbon decomposition and the movement of bacterial cells has a large impact on regulating soil biogeochemical cycles^1,2^. In addition, the transport of pathogenic bacteria from fecal waste to drinking water reservoirs poses risks to human health^1^. Furthermore, many soil bioremediation projects rely on the injection of contaminant-degrading bacteria such as petroleum-degrading *Bacillus* sp. to decompose contaminations, and accordingly, the spread of bacteria impacts the remediation efficiency^3–5^. Mechanistic understanding of how bacterial cells spread in soil is needed to predict soil biogeochemical cycles and improve soil quality, yet such understanding is currently incomplete.

In most current studies, bacterial spreading was attributed to advection and hydrodynamic dispersion^6,7^, filtration, adsorption, and desorption^8^, as well as bacterial motility^9^. Many studies also show that surfactants can facilitate bacterial transport in porous media^10,11^. In addition to the above mechanisms, a recent study^12^ discovered a new bacterial spreading mechanism that *Pseudomonas aeruginosa*, a major human pathogen and bacterium found in soil, can self-generate flows along sharp corners and spread in a synthetic soil by producing biosurfactants that change the wettability of solid surfaces.

In this study, we investigate whether biosurfactant-enabled bacterial spreading is a conserved mechanism of soil bacteria. We hypothesize that biosurfactant-based bacterial spreading mechanism is common in soil because many soil bacteria produce surfactants, including *Bacillus subtilis*^13,14^, *Pseudomonas fluorescens*^15,16^, *Pseudomonas putida*^17,18^. This study focuses on *B. subtilis* and *P. fluorescens*, two species of plant growth-promoting rhizobacteria^19^ that are ubiquitous in soil^20,21^. The surfactin produced by *B. subtilis* and the rhamnolipid synthesized by *P. fluorescens* are the most analyzed biosurfactants due to their application in bioremediation of diesel-contaminated soil^22^ as well as their biodegradability and low toxicity^23^, and the surfactin is also known for antimicrobial and antifungal activities^24–26^.

Here, we show that surfactant-producing bacteria, *B. subtilis* and *P. fluorescens*, can generate flows along sharp corners of a transparent chamber made from polydimethylsiloxane (PDMS), which has similar surface properties as hydrocarbon-covered hydrophobic soil. We further show that corner flow is surfactant dependent and the contact angle of the solution dictates the critical corner angle for the corner flow. Finally, we show that the maximum height of corner flows generated by bacteria is controlled by the corner geometry, i.e., the roundness or cutoff of the corner, and can be predicted by the capillary theory developed for pure and homogeneous wetting liquids.

## Results

### Corner Flows at Sharp 30° Corners generated by *B. subtilis* and *P. fluorescens*

First, we investigate whether similar to *Pseudomonas aeruginosa*^27^, typical surfactant-producing soil bacteria *Bacillus subtilis* and *Pseudomonas fluorescens* can generate corner flows. We consider three strains: *Bacillus subtilis* 3610 (Wild-type and surfactant-producing strain), *Bacillus subtilis* DS1122 (surfactant-deficient mutant) and *Pseudomonas fluorescens* PF15 (surfactant-producing strain). To test whether these strains generate corner flows, we grow bacteria in M9 solution in a transparent polydimethylsiloxane (PDMS) chamber with four corners, 30°, 60°, 90° and 120° (as shown in Fig. 1A). Afterward, we visualized the spread of the culture medium stained with fluorescein along these four corners using a digital camera over a 24-hour period. Fig. 1B shows a presentative image of the bacterial solution in the chamber at the time of inoculation (*t* = 0 h). As the bacteria grew in the chamber over time, we observed that surfactant-producing bacteria *B. subtilis* 3610 (WT) and *P. fluorescens* PF15 generated corner flows at the 30° corners, as shown in the time-lapse images of the bacterial solution at the 30° corners in Fig. 2. The existence of corner flows at other corner angles and the critical corner angle to generate corner flows will be discussed in the following sections. In contrast to these two surfactant-producing strains, surfactant-deficient mutant *B. subtilis* DS1122 (defective in surfactin), did not generate flow at the 30° corners. These observations confirmed our hypotheses that surfactant-producing bacteria, such as *B. subtilis* 3610 (WT) and *P. fluorescens* PF15, can generate flows along sharp corners by producing surfactants, while surfactant-deficient bacteria like *B. subtilis* DS1122 cannot.

**FIG 1.**
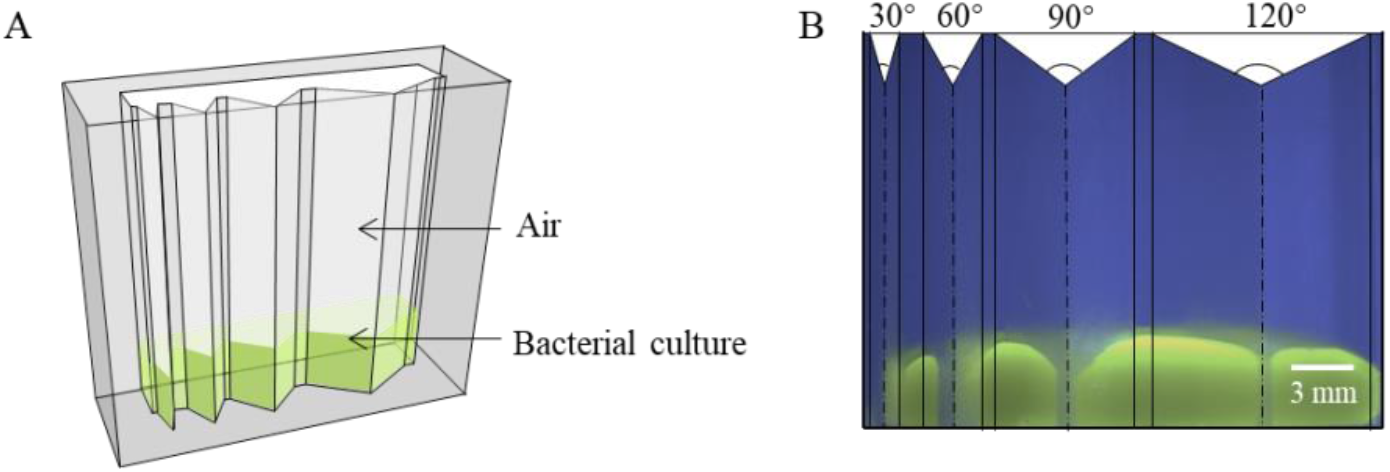
(A) Schematic of experimental corner flow experiments. Bacterial solution (green) was placed in the PDMS chamber with four corner angles. The green color is due to the addition of 0.005% (w/v) fluorescein sodium salt for visualization purpose. The corner angles are 30°, 60°, 90° and 120° from left to right. (B) Image of the bacterial solution in the chamber at *t* = 0 h, which is defined as the time when the bacterial solution was transferred into the chamber.

**FIG 2.**
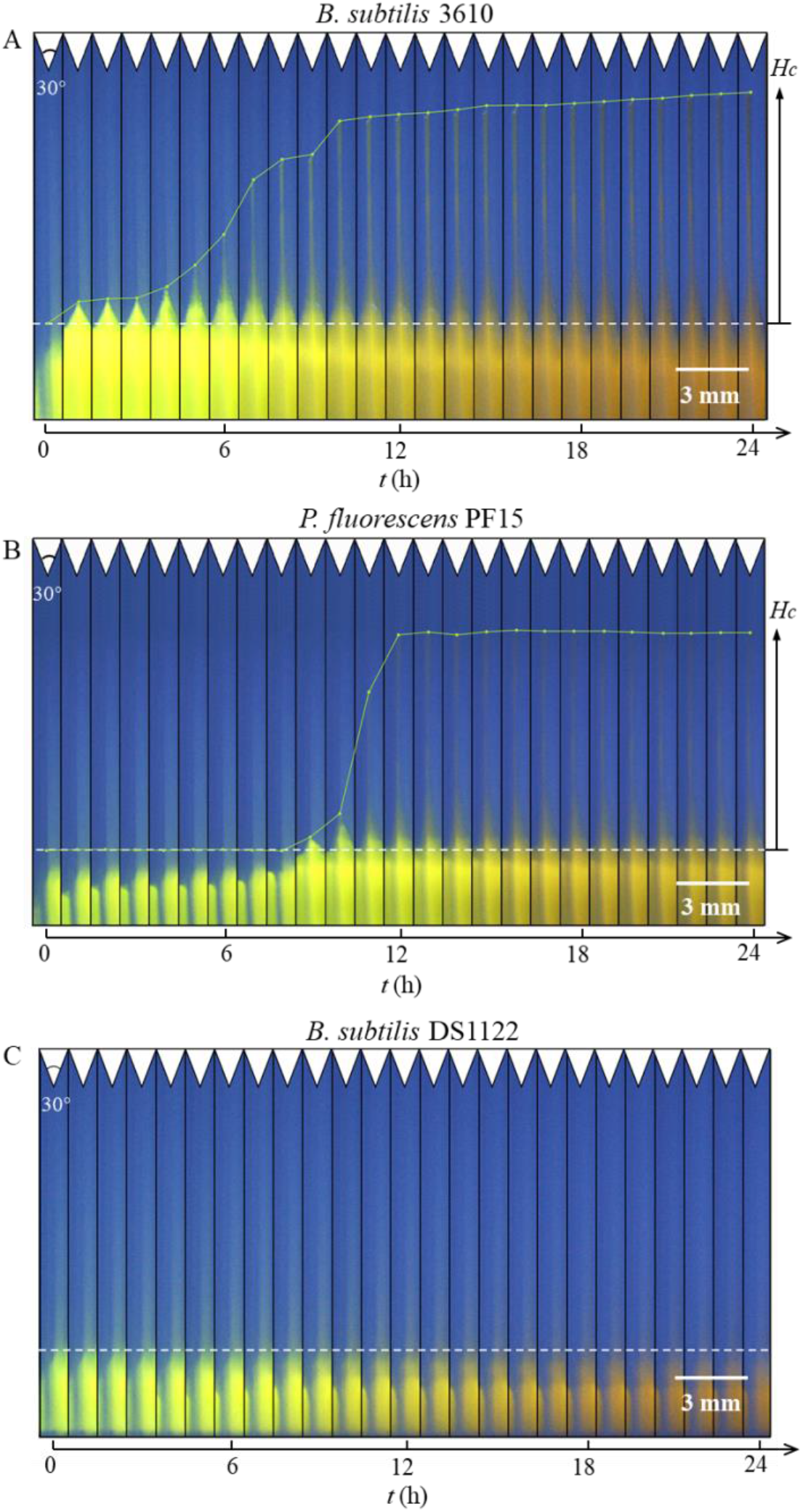
Time-lapse images of corner flow at the 30° corners during a 24-hour bacterial growth period for (A) surfactant-producing strain *B. subtilis* 3610, (B) surfactant-producing strain *P. fluorescens* PF15, and (C) surfactant-deficient strain *B. subtilis* DS1122. Images were cropped at the 30° corner from time sequence images of the chamber with 4 different angles (30°, 60°, 90° and 120°) shown in Fig. 1B. The white dotted horizontal lines represent the initial height of bacterial culture in the chamber at growth time *t* = 0 h. The green lines were added to the image to indicate the tip positions of the corner flows over time. The green color of the bacterial solution is from the 0.005% fluorescein sodium salt added to the bacterial culture. The color of bacterial culture in the chamber gradually turned from bright green to dark yellow due to the increase in bacteria cell density which makes the solution turbid. Note that the contrast and brightness of the figures have been enhanced to increase the visibility of corner flow. The videos of corner flow development with original color are shown in Movie S1, Movie S2 and Movie S3.

Second, we investigate the speed of corner flows at the sharp 30° corner generated by the surfactant-producing bacteria. We plotted the heights of the tip of the corner flows versus time for these three strains shown in Fig. 3. For *B. subtilis* 3610, the corner flow started at *t* = 2 h and ended at *t* = 18 h with a maximum height of 9 mm. For *P. fluorescens* PF15, the corner flow started at *t* = 8 h and ended at *t* = 15 h with a similar maximum height, about 9 mm. The average climb speeds were 0.6 mm/h and 1.3 mm/h for *B. subtilis* and *P. fluorescens*, respectively. The speed of the bacterial corner flow, on the order of mm/h, is similar to bacterial surface swarming, the fastest mode of known bacterial surface translocation^28^.

**FIG 3.**
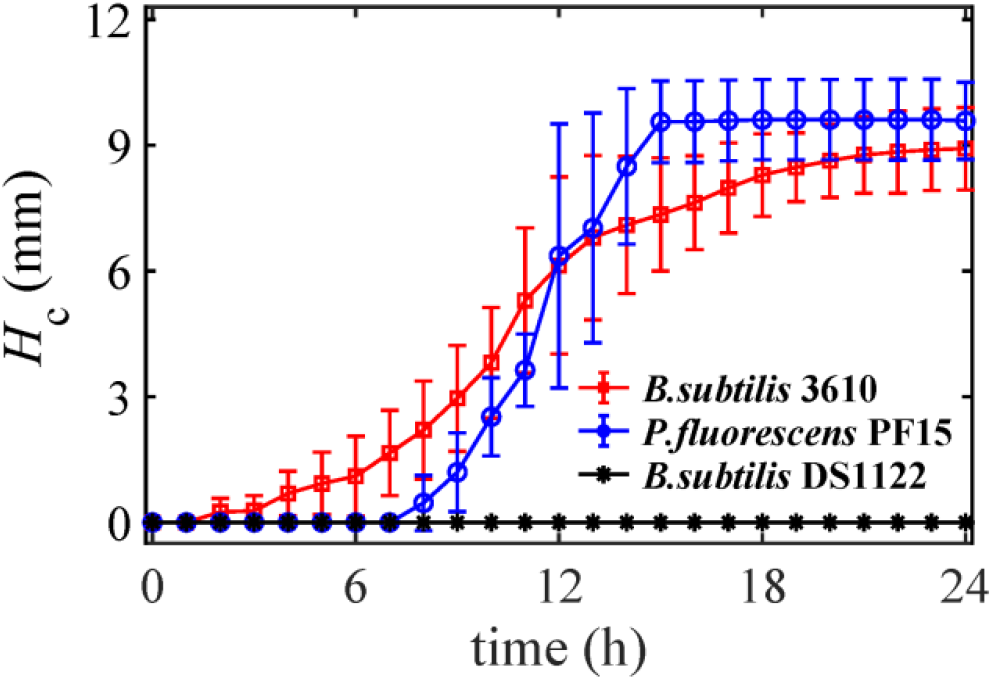
The time evolution of the tip position of corner flows at the 30° corner for *B. subtilis* 3610 (WT), *B. subtilis* DS1122 (defective in surfactin) and *P. fluorescens* PF15. *H*_c_ indicates the tip position of the corner flow from its initial position (shown in Fig. 2). The error bars represent the standard error of measurements of six to seven replicates experiments for each strain.

Further, we show that bacterial cells transport with the bacteria-generated corner flows. We sampled the *B. subtilis* 3610 solution at the tip of the corner flow after 24 hours and diluted it with abiotic M9 solution by about 50 times. Then, we imaged the bacterial sample under a Nikon C2 plus Confocal Laser Scanning Microscope. As shown in Fig. S2, *B. subtilis* 3610 cells exist in the tip of the corner flow, suggesting that surfactant-producing bacteria can indeed make use of the corner flow mechanism to spread.

### Contact angle and the critical corner angle for bacterial corner flows

To demonstrate how bacteria-produced biosurfactants generate corner flows, we measured the optical density OD_600_ and surfactant-related properties, including surface tension (*γ*) and contact angle (*θ*_c_), of the bacterial solution over time. Note that because the volume of the bacterial solution in the PDMS chamber was not sufficient for these measurements, we used bacterial solution grown in 50-mL tubes under identical oxygen, nutrient, and temperature conditions as in the PDMS chamber for measurements. For *B. subtilis* 3610 and *P. fluorescens* PF15, *θ*_c_ and *γ* decreased gradually beginning at *t* = 4 h and *t* = 6 h, respectively. At the beginning of experiments *t* = 0 h, the contact angle of the bacterial solution on the PDMS surface was *θ*_c_ ≈ 115° for all strains, because the solution was similar to water, whose contact angle on PDMS is about 117°^29^. At *t* = 16 h, the wettability of PDMS for both strains, i.e., *B. subtilis* 3610 and *P. fluorescens* PF15, changed from initially non-wetting (*θ*_c_ ≈ 115°) to wetting (*θ*_c_ ≈ 60°). In comparison, for the surfactant-deficient strain *B. subtilis* DS1122, despite a slight decrease in surface tension *γ*, the contact angle *θ*_c_ of the bacterial solution on the solid surface remained above 90°, thus the surface remained non-wetting. For the surfactant-producing strain *B. subtilis* 3610, *θ*_c_ dropped from 115° (nonwetting) to 60° (wetting) during *t* = 4 to 17 hours (Fig. 5A), which started earlier than *P. fluorescens*, for which *θ*_c_ dropped during *t* = 6 to 17 hours (Fig. 5B). Consistently, as shown in Fig. 3, the beginning time of corner flows in chambers for *B. subtilis* 3610 was around *t* = 2 h, earlier than the start time of the corner flow of *P. fluorescens*, which is around *t* = 7 h. The start time of corner flow and the time when the contact angle starts to change for *B. subtilis* and *P. fluorescens* are similar, suggesting that the bacterial corner flow was indeed caused by the surfactant-induced change in the contact angle of the solution on surfaces.

Next, we show that the critical corner angle for bacteria to generate corner flow can be predicted from the contact angle of bacterial solution on surfaces. According to classical corner flow theory, e.g., Concus–Finn criterion^30^, corner flow occurs when 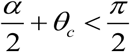 (where *α* is the corner angle and *θ*_*c*_ is contact angles). For *B. subtilis* 3610, after bacteria produced sufficient surfactants, the advancing contact angle on the PDMS surface reduced to around 60° as shown in Fig. 4A, so the predicted critical corner for corner flow is *α*_*th*_ = 2 × (*π* / 2 − *θ*_*min*_) = 2 × (90° − 60°) = 60°. We prepared PDMS chambers which contain interior corners of different degrees of 30°, 50°, 60°, 90° and 120°. And we transferred 600 – 800 *μ*L bacterial culture at OD_600_ between 0.5 ± 0.1 into a chamber and imaged the position of the bacterial solution over 24 hours. During the experiments, we observed noticeable corner flows along corners with 30° and 50° corners, but no fluid rises along 90° or 120° corners. As for 60°corners, a slight rise occurs. These experimental results suggest that the critical corner angle for *B. subtilis* is about 60°, which is consistent with the predicted critical corner angle *α*_*th*_ = 60° calculated by classic corner flow theory. For *P. fluorescens* which also reduced the contact angle to 60°, so the predicted critical corner angle is also *α*_*th*_ = 60°. Our experiments in the chamber with 4 different angles (30°, 60°, 90° and 120°) (see *Supporting Information* Video) also show corner flows at 30° corner but no fluid rise in 90° or 120° corners. A slight rise at the 60° corner was also observed, indicating the critical corner angle for *P. fluorescens* is also about 60°. The agreement of the predicted and the observed critical corner angle for both surfactant-producing bacteria, *B. subtilis* and *P. fluorescens*, further confirms that the bacterial corner flow is due to and can be predicted by biosurfactant-induced changes in contact angle.

**FIG 4.**
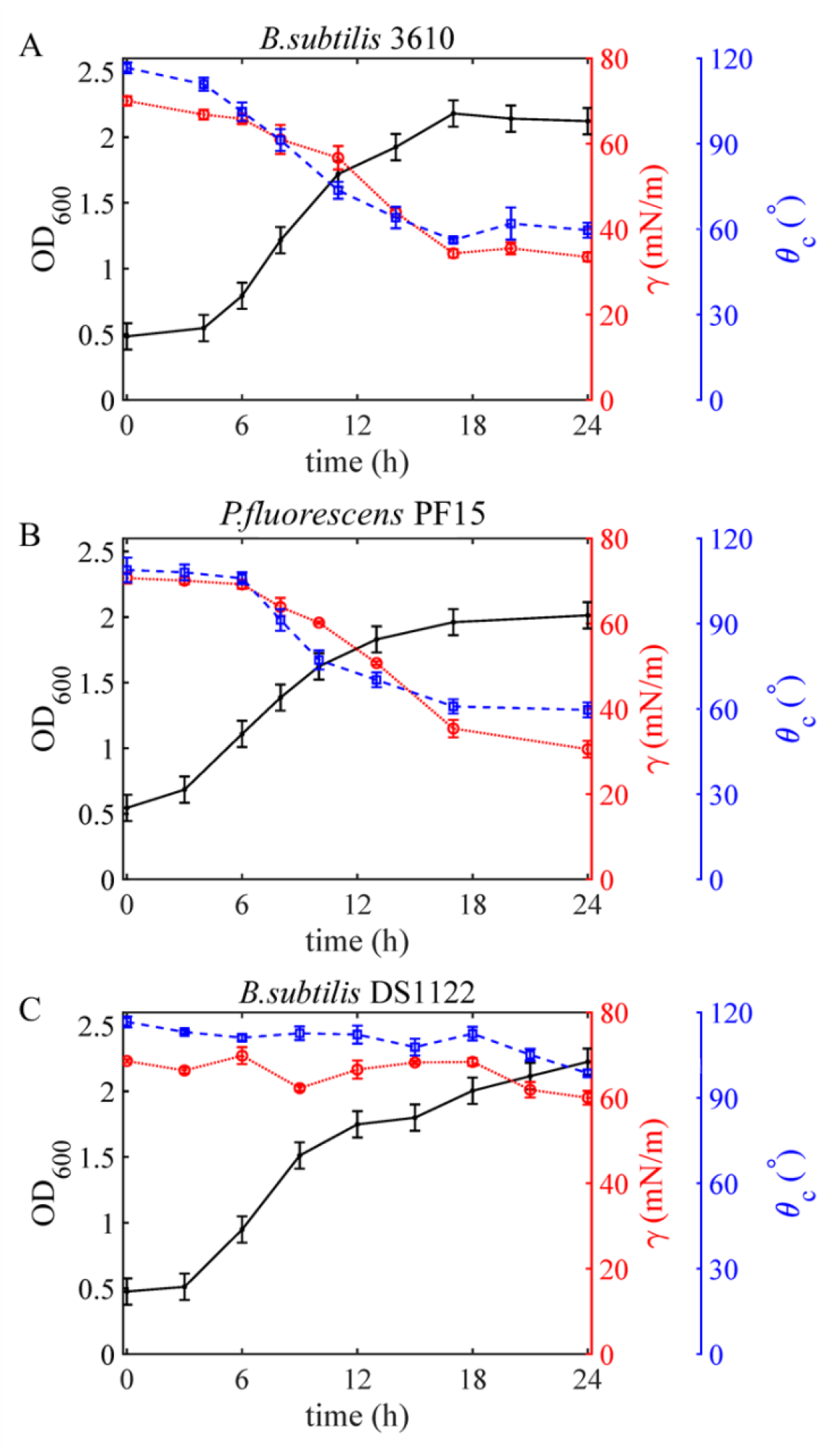
The time evolution of the advancing dynamic contact angle *θ*_c_ on the PDMS surface, the surface tension *γ*, and the cell density OD_600_ of bacterial solutions for *B. subtilis* 3610 (WT), *B. subtilis* DS1122 (defective in surfactin) and *P. fluorescens* PF15, separately. The error bars represent the standard error of measurements of 3 to 4 liquid drops.

**FIG 5.**
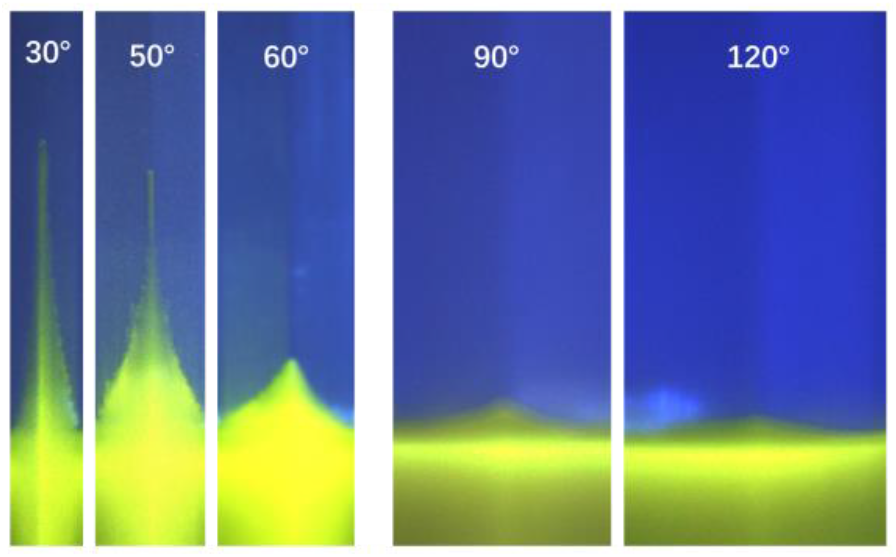
Image of solutions containing *B. subtilis* 3610 cells along with five different corners after 20-hour growth period. The initial bacteria density was OD_600_ ≈ 0.5 ± 0.1. The first three images with corners of 30°, 50°, 60° were cropped from one chamber (see *Supporting Information*), and the last two images with corners of 90°, 120° were cropped from the chamber shown in Fig. 1.

### Surfactant concentration and corner flows

Next, we investigate the concentration of surfactants produced by bacteria that can generate corner flows. Specifically, we repeat the corner flow experiments using various concentrations of commercially-available surfactants rhamnolipid (Sigma) and surfactin (Sigma), which are the surfactants produced by *P. fluorescens*^31,32^ and *B. subtilis*^33,34^, respectively. Here, we measured the surface tension and contact angle of surfactant solutions at different concentrations and transferred surfactant solutions into chambers to observe the rise of corner flow at the 30° corner. As the surfactant concentration increases, the contact angle of the liquid decreases and corner flows start to occur. As shown in Fig. 6, the surface tension *γ* and contact angle *θ*_c_ for surfactin and rhamnolipid solutions dropped rapidly when the concentration was in the range of 1 × 10^−5^ - 1 × 10^−4^ M and 1 × 10^−5^ - 1 × 10^−2^ M, respectively. *γ* and *θ*_c_ were determined from the shape of pedant droplet and the angle of moving droplet as shown in the insets of Fig. 6 (see Methods for details). Corner flows were only observed when the surfactant concentration was higher than the critical value, which for surfactin is about 2 × 10^−5^ - 3 × 10^−5^ M and for rhamnolipid is about 1 × 10^−4^ - 3 × 10^−4^ M.

**FIG 6.**
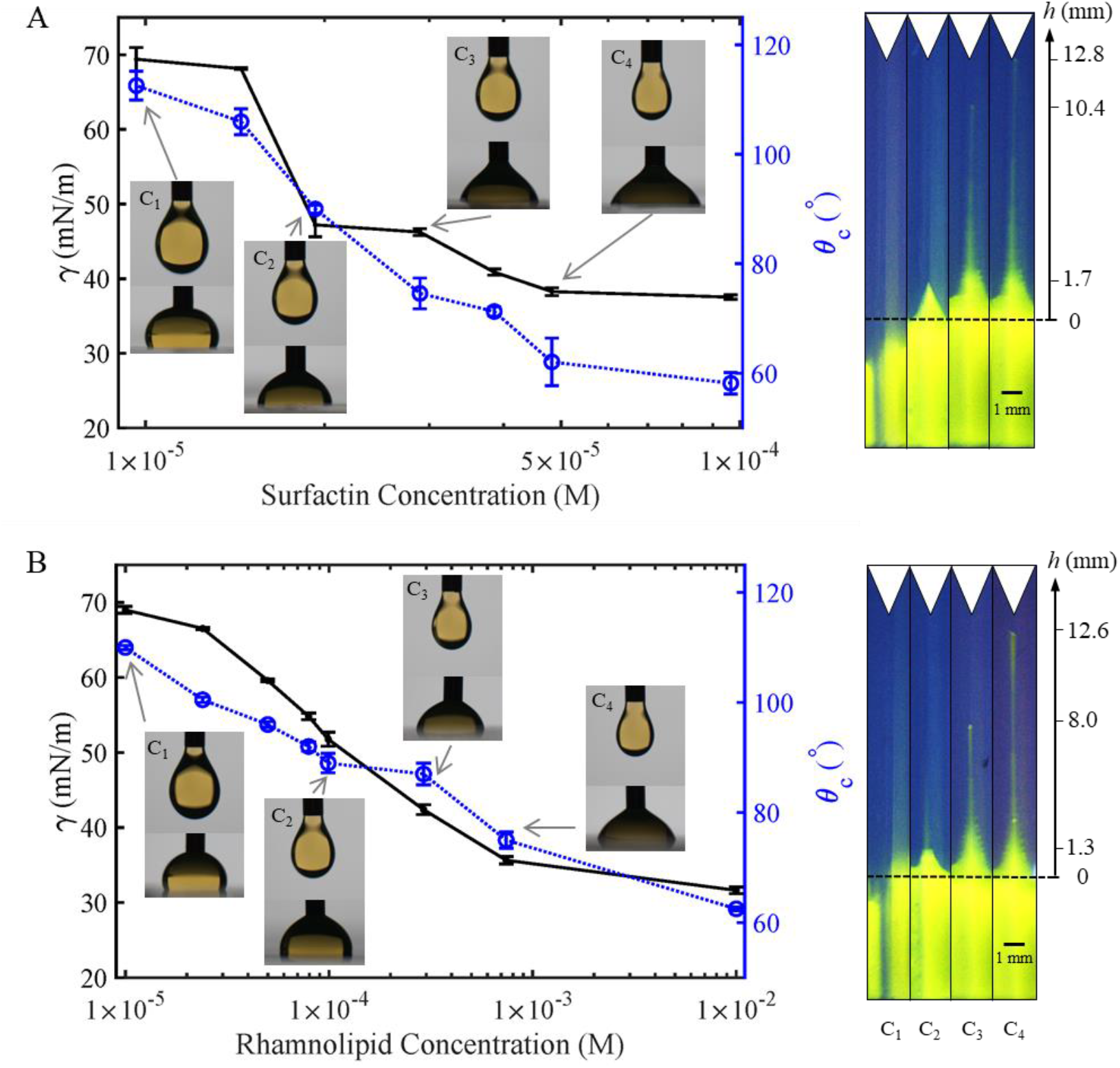
The surface tension *γ* and contact angle *θ*_c_ for pure surfactin solution (A) and rhamnolipid solution (B) at different concentrations. The insets show examples of pedant droplet used to measure surface tension and moving droplets to measure contact angle (see Methods for details). The right image shows the corner flow generated by the surfactants at different surfactant concentrations (C_1_, C_2_, C_3_, C_4_) indicated on the left figures.

Note that both the corner flows generated by pure bacterial surfactants reached above 12 mm, which is higher than the height of corner flow (9 mm) generated by bacteria culture. We hypothesize that the smaller corner height for bacterial solution is caused by the evaporation of bacteria solution during 24-hour growth period.

### Maximum height of corner flow

During our experiments, we observed that tip of corner flows generated by *B. subtilis* 3610 and *P. fluorescens* PF15 at the 30° corner stopped reached a maximum height, *h*_*max*_ ≈ 9 mm, as shown in Fig. 3. However, corner flows can rise to infinite height theoretically if the corner is perfectly sharp^35,36^. We hypothesize that the maximum corner flow height is related to the roundness or cutoff of the corner, because corners can hardly be perfectly sharp^37,38^. The microscopic image of the 30° corner (Fig. 7B) shows that the corner in our experiments, due to the resolution of our 3D printing (see *Methods* for details), is rounded with step-like structures along its inner surface, as shown in Fig. 7C.

**FIG 7.**
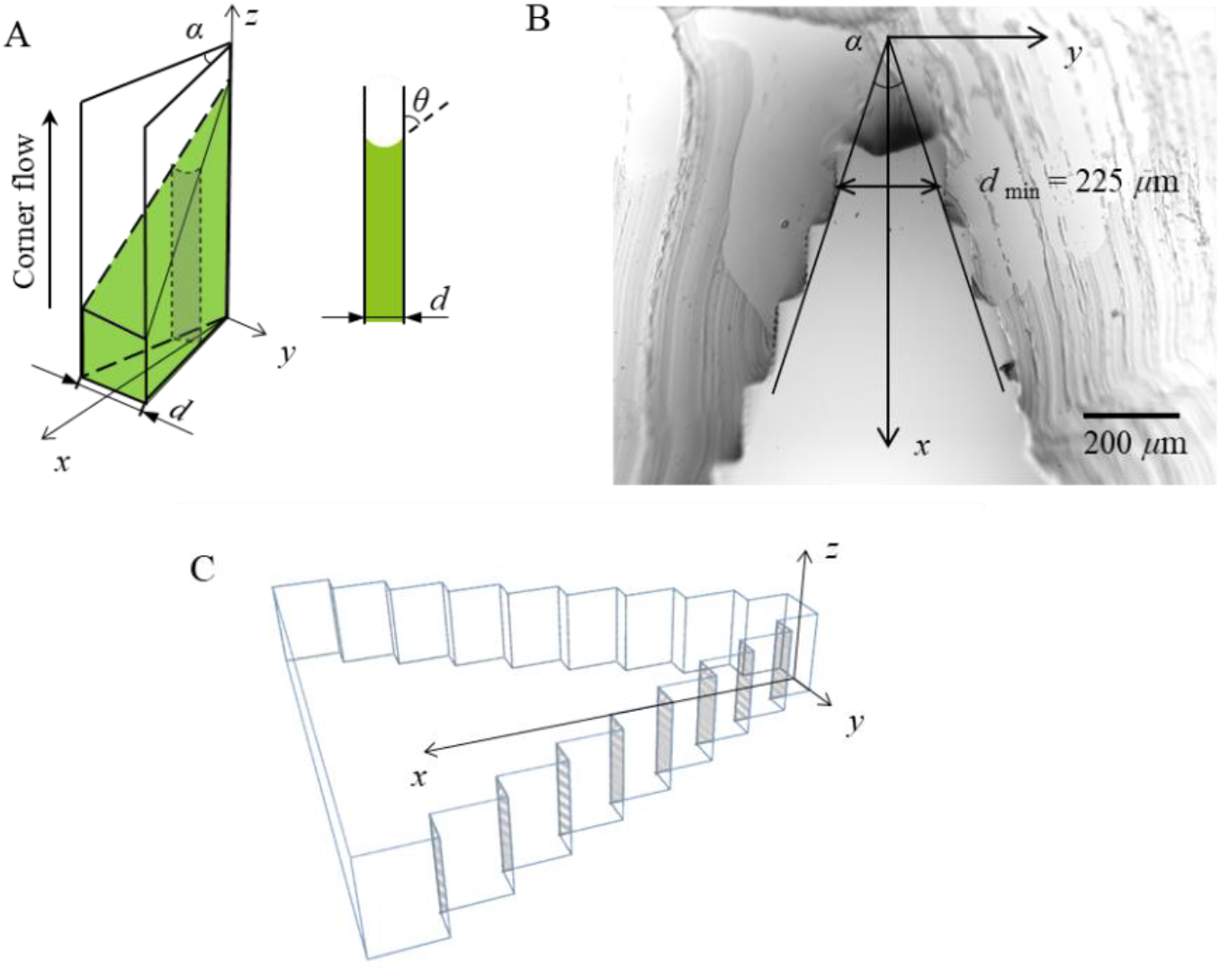
(A) A schematic diagram of the upward flow of the bacterial solution (green) along a 30° corner. A cross-sectional image parallel to the (*y, z*)-plane is shown at the right. *θ*c is the contact angle, *α* is the corner angle, *d* is the local separation changing with the distance *x*. (B) Confocal image of the cross-section of the 30° corner of the PDMS chamber, the minimum separation in the tip position is 225 *μ*m. (C) A schematic of the step-like structures of the 30° corner.

**FIG 8.**
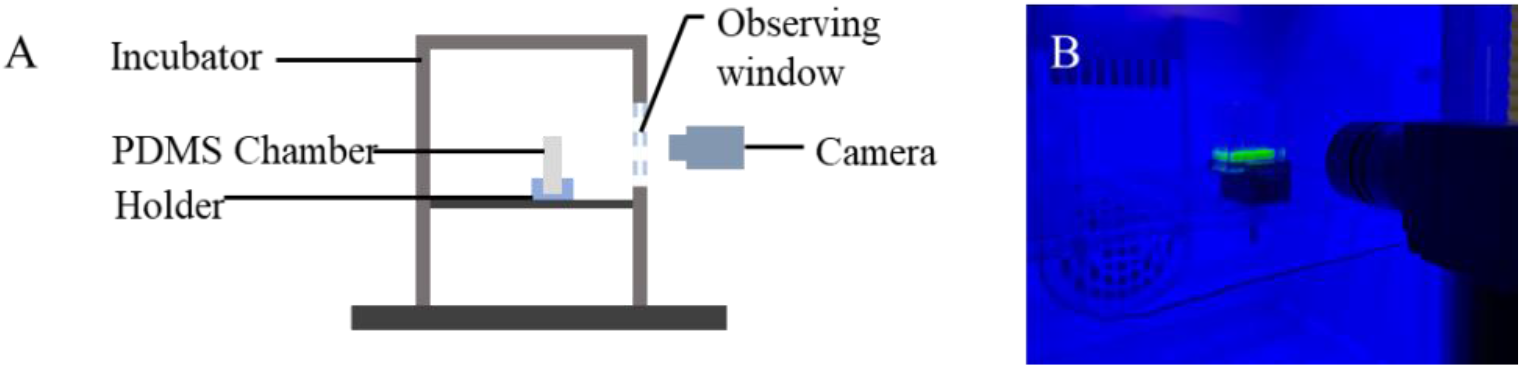
(A) A schematic diagram of the bacterial corner flow experimental setup. (B) An image of the experimental setup for the bacterial corner flow experiments.

To estimate the maximum height of corner flow along the rounded corner, here we use a simple fluid mechanics theory developed for pure wetting liquid^39,40^. The following assumptions were made^41,42^: (i) the motion of the fluid is mainly vertical and dominated by the curvature in the (*y, z*) - plane and the constant capillary pressure can be calculated from the contact angle *θ*_*c*_; and (ii) evaporation of liquid, friction, and inertial effects are not considered. Imagine that the two plates that form the corner are composed of many parallel plates with different distances *d* apart, e.g., consists of step-like structures in the inner surface of the chamber (Fig. 7C). The fluid meets the container wall with a prescribed contact angle *θ*_*c*_. The weight of liquid that rises vertical distance *h* by a segment of the wall off length *l* between two parallel walls with distance *d*, is *F*_*g*_ = *ρgdhl*, where *g* is the free-fall acceleration, *ρ* is the liquid density. The capillary force due to surface tension of the liquid between two parallel plates can be derived using Young-Laplace equation^43^, *F*_*σ*_ = 2*lγ*cos*θ*_c_. Due to force balance, the capillary driving forces (*F*_*σ*_) equal to the gravitational force (*F*_*g*_), thus the maximum height of the liquid rise is:

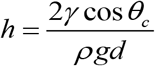

From the microscopic image of the corner (Fig. 7B), the minimum width at the tip of the corner is *d* = 225 *μ*m, which is the minimum width of a series of parallel plates. Substituting *θ*_c_ = 60°, *γ* = 30 mN/m, *ρ* = 1000 kg/m^3^ and *d* = 225 *μ*m into above equation, we found *h* = 13.6 mm. Consistently, our experiment results show that the maximum height of the corner flow at 30° corner is 9 mm for the two surfactant-producing bacteria considered here, which is the same order of magnitude as our theoretical derivation, suggesting that the maximum height of corner flow is indeed limited by the roundness of the corner.

## Discussion

We demonstrate that typical biosurfactant-producing soil bacteria, Wild-Type *Bacillus subtilis* and *Pseudomonas fluorescens*, can self-generate flows along sharp corners with corner angle less than 60°. We show that a surfactant-deficient mutant of *Bacillus subtilis* did not generate the corner flow, thus bacterial motility is not needed for bacteria to generate corner flows. The speed of corner flows shown is on the order of millimeters per hour, similar to bacteria swarming, the fastest mode of known bacterial surface translocation^44,45^. Further, we show that the bacterial corner flow was generated by surfactant-induced change in contact angle of the bacterial solution on the solid surface, and the critical corner angle to generate corner flow can be predicted by the contact angle. Finally, we demonstrate that the maximum height of bacterial corner flow can be predicted by the corner geometry, i.e., the roundness or cutoff length of the corners. We anticipate that the bacterial corner flow revealed in this study is prevalent in soil where biosurfactant-producing bacteria^46,47^ and angular pores^48–50^ are common^51^. Our results also suggest that both bacteria-produced surfactants and the geometry of the soil pore network should be considered in predicting biosurfactant-induced corner flows and bacterial spreading. The mathematical description of biosurfactant-driven flow developed in this study will help improve predictions of bacteria transport in soil and facilitate designs of soil-based bioremediation projects.

## Materials and Methods

### Bacterial Strains and Growth

The bacterial strains used in the present study were *Bacillus subtilis* 3610 (Wild-type), *Bacillus subtilis* DS1122 and *Pseudomonas fluorescens* PF15. Strain cells were streaked from - 80°C freezer stocks onto an LB Medium plate (1.5% agar). *P. fluorescens* and *B. subtilis* were grown at 30°C or 37°C, respectively.

### Bacterial Solution

5 mL of LB liquid medium were inoculated with cells from an isolated colony on the plate into a 50-mL tube. The LB medium inoculated with *B. subtilis* was incubated on a shaker at 37°C, 200 rpm, and the LB medium inoculated with *P. fluorescens* was placed on a shaker at 30°C, 200 rpm overnight. The bacterial overnight cultures were subjected to centrifugation at 4,000 rpm for 10 minutes. Afterward, we removed the supernatant and diluted the bacterial cells at the bottom of the tube with M9 medium and mixed them using a vortex mixer. The cell density of culture was diluted to OD_600_ = 0.5 ± 0.1 by tuning the volume of M9 medium. The M9 medium used in this study was supplemented with 0.03 *μ*M (NH_4_)_6_(Mo_7_)_24_·4H_2_O, 4 *μ*M H_3_BO_3_, 0.3 *μ*M CoCl_2_·6H_2_O, 0.1 *μ*M CuSO_4_·5H_2_O, 0.8 *μ*M MnCl_2_·4H_2_O, 0.1 *μ*M ZnSO_4_·7H_2_O, 0.1 *μ*M FeSO_4_·7H_2_O and 2% glucose. When noted, 0.005% (w/v) fluorescein sodium salt was added.

### 3D-Printed Molds Preparation and Fabrication of the PDMS Slabs

We use 3D-printed molds to produce the PDMS slabs used in the experiment. The mold is composed of a cuboid (30 mm × 25mm × 4 mm) and four triangular prisms (the image of the mold is shown in *Supporting Information*). The heights of the cross-sections of these triangular prisms are all 3 mm. The mold with four different angles was printed by a 3D printer (Anycubic Photon Mono X) using a 405nm UV resin (Anycubic). The printed molds cannot be directly used for PDMS casting because chemicals released from 3D-printed objects will inhibit PDMS curing in the vicinity of these objects^52^. So before casting, the printed mold was UV post-curing for 20 minutes, immerged in isopropanol for 6 hours, then treated with air plasma corona (BD-20AC) for 1 min, and then silanized using triethoxy (1H, 1H, 2H, 2H-perfluoro-1-octyl) silane for 3 h. Then the mold was transferred to a petri dish and a 10:1 w/w base/curing agent PDMS liquid was poured onto the 3D printed mold. The composite was cured in a hotplate at 80 °C for at least 2 hours.

### Corner Flow Experiment

For the corner flows experiment, a sterilized PDMS chamber was placed in an incubator with a transparent front door. We transferred 600 – 800 *μ*L prepared bacterial culture at OD_600_ = 0.5 ± 0.1 into the PDMS chamber using a 3-mL syringe. The incubator was set to the temperature of 37 ± 2 °C for *B. subtilis* and 30 ± 2°C for *P. fluorescens*, and relative humidity was kept at 80 ± 10%. To visualize the bacterial-induced corner flow, a blue LED light was placed in the incubator and the M9 medium containing fluorescent 2-NBDG glucose to make the slim corner flow visible. A digital camera (Blackfly S BFS USB3, Teledyne FLIR) was placed in front of the chamber and set to take photos at 2 mins intervals for 24 hours. The length of corner flow was measured from the initial reservoir level to the top of corner flow.

### Contact Angle and Surface Tension Measurements

To quantify the biosurfactant related parameters, we measured the time evolution of contact angle *θ*_c_, surface tension γ, and bacterial cell density (OD_600_) over time for 24 hours. Due to the limited volume of bacterial solution in the PDMS chamber, we grow bacterial solution in tubes to mimic the growth of bacteria in chambers. We diluted overnight bacterial culture to OD_600_ = 0.5 ± 0.1 and separated the uniform culture by transferring 5 mL aliquots into multiple 50-mL centrifuge tubes. We placed the tubes containing 5 mL cultures (OD_600_ = 0.5 ± 0.1) in a shaking incubator. Specifically, the temperature was set to 37°C for *B. subtilis* and 30°C for *P. fluorescens*. For each data point in Fig. 5, we removed one tube from the incubator and transferred 1 mL bacterial culture to a cuvette and measured the OD_600_. Because we split the 24-hour continuous monitoring into two 12-hour periods and the initial OD_600_ was diluted to 0.5 ± 0.1, the error bar/uncertainty of each OD_600_ data point was set to 0.1.

To measure surface tension *γ* and contact angle *θ*_c_, bacteria culture was centrifuged at 4,000 rpm for 10 minutes and the supernatant was filtered through a 0.2-*μ*m filter to remove bacterial cells. Afterward, we transferred the filtered solution into a 10-mL syringe with an 18-gauge needle. We placed the syringe on a syringe pump connected to a needle and a drop of the solution was pushed out of the needle. To measure surface tension *γ*, the shape of the pendant drops below the needle was recorded and the profile of the droplet edge was fitted using the MATLAB code developed by the Stone group based on the algorithm proposed by Rotenberg et al.^53^. To measure the advancing contact angle *θ*_c_, we injected bacterial solution to the surface of a PDMS slab and imaged the shape of the advancing drop using the syringe pump at a 1.2 mL/hour flow rate. After identifying the edges of the moving drops, we estimated *θ*_c_ as the angle between the PDMS surface and the tangent line of the drop edge near the contact line. Examples of pendant drop and advancing drop are shown in the *Supporting Information*.

## Data Availability

The MATLAB codes for image processing and the estimation of surface tension and contact angle have already shared by J Yang on GitHub: https://github.com/JudyQYang/Bacterial_corner_flow_codes. All other study data were extracted from the Supplementary Movies and are available from the corresponding authors on reasonable requests.

## Acknowledgements

This research was supported by J Yang’s startup fund. L Yuan was supported by the fellowship of Civil, Environmental, and Geo-Engineering at the University of Minnesota. J Sanfilippo was supported by a NIH K22 grant 5K22AI151263-02. We thank Mohamed Donia (Princeton University) for giving us the bacterial strain *Pseudomonas fluorescens* PF15.

